# Plasma Proteome Perturbation for CMV DNAemia in Kidney Transplantation

**DOI:** 10.1101/2022.08.25.505318

**Authors:** Tara K. Sigdel, Patrick Boada, Maggie Kerwin, Priyanka Rashmi, David Gjertson, Maura Rossetti, Swastika Sur, Dane Munar, James Cimino, Richard Ahn, Harry Pickering, Subha Sen, Rajesh Parmar, Benoit Fatou, Hanno Steen, Joanna Schaenman, Suphamai Bunnapradist, Elaine F. Reed, Minnie M. Sarwal, CMV Systems Immunobiology Group

**Affiliations:** Department of Surgery, University of California San Francisco, San Francisco, CA; Department of Pathology and Laboratory Medicine, University of California Los Angeles, Los Angeles, CA; Department of Microbiology and Immunology, University of California Los Angeles, Los Angeles, CA; Department of Medicine, University of California Los Angeles, Los Angeles, CA; Department of Pathology, Harvard Medical School, Boston, MA

## Abstract

**Background:** Cytomegalovirus (CMV) infection, either *de novo* or as reactivation after allotransplantation and chronic immunosuppression, is recognized to cause detrimental alloimmune effects, inclusive of higher susceptibility to graft rejection and substantive impact on chronic graft injury and reduced transplant survival. To obtain further insights into the evolution and pathogenesis of CMV infection in an immunocompromised host we evaluated changes in the circulating host proteome serially, before and after transplantation, and during and after CMV DNA replication (DNAemia), as measured by quantitative polymerase chain reaction (QPCR).

**Methods:** LC-MS-based proteomics was conducted on 168 serially banked plasma samples, from 62 propensity score-matched kidney transplant recipients. Patients were stratified by CMV replication status into 31 with CMV DNAemia and 31 without CMV DNAemia. Patients had blood samples drawn at protocol times of 3- and 12-months post-transplant. Additionally, blood samples were also drawn before and 1 week and 1 month after detection of CMV DNAemia. Plasma proteins were analyzed using an LCMS 8060 triple quadrupole mass spectrometer. Further, public transcriptomic data on time matched PBMCs samples from the same patients was utilized to evaluate integrative pathways. Data analysis was conducted using R and Limma.

**Results:** Samples were segregated based on their proteomic profiles with respect to their CMV Dnaemia status. A subset of 17 plasma proteins was observed to predict the onset of CMV at 3 months post-transplant enriching platelet degranulation (FDR, 4.83E-06), acute inflammatory response (FDR, 0.0018), blood coagulation (FDR, 0.0018) pathways. An increase in many immune complex proteins were observed at CMV infection. Prior to DNAemia the plasma proteome showed changes in the anti-inflammatory adipokine vaspin (SERPINA12), copper binding protein ceruloplasmin (CP), complement activation (FDR=0.03), and proteins enriched in the humoral (FDR=0.01) and innate immune responses (FDR=0.01).

**Conclusion:** Plasma proteomic and transcriptional perturbations impacting humoral and innate immune pathways are observed during CMV infection and provide biomarkers for CMV disease prediction and resolution. Further studies to understand the clinical impact of these pathways can help in the formulation of different types and duration of anti-viral therapies for management of CMV infection in the immunocompromised host.

## Introduction

Cytomegalovirus (CMV) infection is a frequent infectious complication in an immunocompromised host, as is the case in an organ transplant recipient who faces life-long immunosuppression^1^. CMV infection can occur either from reactivation of recipient or donor CMV or *de novo* infection in the recipient after transplantation^2^. The clinical spectrum ranges from asymptomatic to systemic infection and end-organ disease with pneumonitis, meningitis, esophagitis, and colitis. Furthermore, CMV infection also results in an increased host susceptibility to secondary infections and increased alloimmunity, leading to acute and chronic allograft rejection^3^. Evaluation of peripheral mRNA changes from clinical and sub-clinical CMV primary infection and reactivation has provided insight into causal factors for allograft dysfunction secondary to CMV^4-6^, host biological pathways actively altered by the virus^7^, and the dynamic temporal changes of cell cycle, DNA damage molecules^8-12^ that occur during the evolution of CMV viral replication^13^.

Mass spectrometry-based proteomics of human serum and plasma has been used for identifying markers for disease diagnosis^14^, and for the study of viral infections such as influenza^15^, COVID-19^16,17^, and HIV^18^. Characterizing the host response to CMV infection will allow for an increased understanding biological events associated with CMV infection severity and will also provide insights into the negative impact of CMV infection on the transplanted organ. CMV infection drives alloimmune injury of the engrafted organ through heterologous immunity in an organ transplant recipient and is an important confounder for higher rates of graft failure after kidney transplantation^19,20^. To address these key questions, we have utilized a highly characterized, propensity-matched cohort of kidney transplant patients with and without post-transplant CMV DNAemia, with serially collected samples at 3 months (pre-CMV infection) and 12 months post-transplant (post-CMV infection), as well as immediate post-CMV time points at 1 week and 1 month post CMV DNAemia and also utilized preexisting transcriptomic data on time-matching PBMCs from the same cohort.

## Materials and Methods

### Patient characteristics

Peripheral blood samples were collected from kidney transplant recipients enrolled in an IRB-approved study from the University of California Los Angeles and plasma samples were isolated^21^. Patients received induction with either anti-thymocyte globulin (ATG) or basiliximab depending on pre-transplant levels of sensitization and donor kidney quality followed by protocolized immunosuppression with tacrolimus, mycophenolate mofetil, and prednisone, as previously described^22^. CMV prevention was performed according to center protocols summarized as follows: 6 months of valganciclovir for high-risk donor positive (D+) and recipient negative (R-) patients and 3 months of valganciclovir for intermediate risk recipient positive (R+) patients who received ATG induction. R+ patients who received basiliximab or low risk (D-/R-) patients received acyclovir prophylaxis to prevent HSV and VZV infection. All study subjects underwent regular CMV PCR screening at 3-, 6- and 12-months post-transplantation to detect CMV DNA in peripheral blood, and further at 1 week and 1 month after detection of CMV viremia.

### Patient selection

A matched cohort of patients with (n=31) and without (n=31) CMV DNAemia post-transplant (>137 IU/ml), were identified by review of clinical records, and the following blood samples were processed for proteomics: (1) Baseline (3-month post-transplant sample, prior to DNAemia start) (2) One-week post-DNAemia (week 1, early post-CMV) (3) One-month post-DNAemia (month 1, intermediate post-CMV) (4) 12 month after transplantation (long-term post CMV). Four of the 31 patients that developed CMV DNAemia after transplantation were donor IgG seropositive, and recipient seronegative; the others were recipient IgG seropositive. For patients with multiple episodes of CMV DNAemia over 137 IU/ml, the first episode closest to transplantation was studied. Patients with a history of CMV DNAemia were matched on a 1:1 basis to a cohort of kidney transplant recipients without a history of CMV DNAemia via propensity scores estimated from a logit model with variables donor type (deceased versus living donor status), recipient, sex, race, and induction type in kidney transplant recipients who were either D+/R-or R+. Samples were selected for each control patient that corresponded in terms of time post-transplant with the baseline and long-term CMV DNAemia patient samples. Of these CMV DNAemia negative patients, 24 were CMV seropositive and 7 were seronegative with CMV positive donors.

### Targeted LC/MS-based proteomics

The plasma samples were processed using a proteomics blotting methodology in a 96-well format^4,23^. In brief, 1 µL of plasma (∼50 µg of protein) was added to 100 µL of urea buffer (8 M urea in 50 mM ammonium bicarbonate buffer (ABC). Following reduction using dithiothreitol (DTT, 50 mM in urea buffer) and alkylation of the cysteine side chains using iodoacetamide (IAA, 250 mM in urea buffer), 10-15 µg of proteins were loaded onto a 96-well plate with an activated and primed polyvinylidene fluoride (PVDF) membrane at the bottom (Millipore-Sigma). Proteins adsorbed to the membrane were trypsinized by incubation with the protease for 2 hours at 37°C. Resulting tryptic peptides were eluted off the membrane with 40% acetonitrile (ACN)/0.1% formic acid (FA). The peptides were subsequently desalted using a 96-well MACROSPIN C18 plate (TARGA, The NestGroup Inc.). Approximately 1 µg of tryptic peptides was analyzed using a Mikros Liquid Chromatography connected to LCMS 8060 triple quadrupole mass spectrometer (both: Shimadzu Corp.). The mass spectrometer was operated in MRM mode. The proteotypic peptide had been selected and validated using pooled plasma samples. A total 629 unique peptides corresponding to 315 proteins were monitored in each LC/MS run with a total run time of 15 minutes per sample. Each peptide was monitored using 3 transitions. The raw data were exported into Skyline software (v20.2.1.315)^24^ for peak area and retention time refinement. The means of the peptide intensities were used for the different protein abundances, which were exported for further analysis.

### Data Analysis

Data preprocessing was conducted with Python (3.8.8), Pandas (1.2.0), Numpy (1.19.5), and Re (2.2.1). The data was set at a 90% limit for missing data. Any aberrated or missing values were imputed by the minimum value observed in a similar cohort member. Moreover, all raw abundance data was log2 normalized prior to any further analysis. All exploratory data analysis and descriptive statistical plots were generated by using Seaborn (0.11.1). Differential Expression Analysis was conducted by linear models by using R (4.0.4), limma (3.46.0) and ggplot2 (3.3.5). Three major comparisons were created to discern a CMV + signature: CMV+ Baseline samples vs all CMV-samples, CMV+ samples vs all CMV-samples, and CMV+ longitudinal samples vs CMV-longitudinal samples. Statistical significance was accepted with a p-value less than or equal to 0.05. Corresponding results were illustrated as volcano plots using ggplot2. Additionally, statistically significant proteins per each time point were summarized in a Venn diagram by using Python (3.8.8) & Matplotlib_venn (0.11.6). Multivariable analysis for clinical confounders revealed no bias for gender, race, ethnicity, age, donor Type, recipient age, induction immunosuppression, or CMV risk group based on recipient and donor CMV IgG status, and time post-transplant for the time of CMV reactivation or infection.

A two-component linear discriminant analysis was used to distinguish the clustering of patient CMV status over time (Python 3.8.8, Sklearn: 0.24.1). Furthermore, a XGBoost machine learning model (Python 3.8.8, XGBoost: 1.4.1) was utilized to identify significant proteins that are associated with CMV DNAemia positivity. The data was normalized by standard scaling. The mean and standard deviation was computed by fit transformation (Sklearn: 0.24.1). Data was then split to preserve 30% data for testing. A param grid of learning_rate, min_child_weight, gamma, subsample, n_estimators, colsample_bytree, and max_depth was automated by using a randomized search cv. The random search was composed of 50 iterations, scoring based on roc_auc, 4 jobs, and a cross-validation composed of a stratified k fold search of 10 splits. Root Mean Square Error (RMSE, 0.345033), Prediction Accuracy (88.1%), CV score (85.21%), and AUC (96%) were used as metrics to observe the accuracy and validity of the model. Moreover, the top contributing features were extracted from the machine learning model and evaluated by the F score; resulting in a set of 37 unique proteins that was run through STRING DB (functional protein association networks) to determine the GO Biological Processes, Molecular Functions, and proteomic interactions.

### Proteome & Transcriptome Correlation

We utilized a previously reported^6^ matching transcriptome data on PBMCs collected from the same cohort with matching time points to perform an integrative multi-omic analysis. Data was downloaded from Gene Expression Omnibus (GSE168598), in which there were 153 overlapping time-matching samples present with an overlap of 186 gene/protein present in both transcriptome and proteome datasets. Transcriptome data was converted into log(cpm) analyzed using R (4.0.4), edgeR (3.32.1) and limma-voom transformation (3.46.0). Differential expression was then calculated for a baseline signature, CMV signature and longitudinal signature. The results were then analyzed for a Pearson correlation between all matching gene transcripts and proteins by using R’s corrplot (0.92).

## Results

### Patient characteristics

A summarization of patient demographics is provided in **Table 1**. As expected, due to the propensity matching design for patient selection, age, sex, race, and induction type were comparable between CMV DNAemia positive (≥ 137 IU/mL of CMV detected in a PCR test: abbreviated PCR+) and CMV DNAemia negative (PCR-) patients (**Table 1**). A similar number of patients were high risk for CMV (D+/R-) as intermediate risk (R+) in both groups. Patients experiencing CMV DNAemia were PCR positive at a median of 80 days after transplantation (range: 10 to 561 days). Median peak viral load was 757 IU/ml (range 146 to 13900 IU/ml). Two CMV DNAemia positive patients were diagnosed with clinical CMV disease, based on standard definitions^25^.

**Table 1.** Demographic and clinical characteristics of kidney transplant recipients with and without detectable CMV viremia.

|  | **CMV DNAemia**<br>**(n=31)** | **No CMV**<br>**DNAemia**<br>**(n=31)** | **p-value** |
| --- | --- | --- | --- |
| **Sex (% Male)** | 68% | 61% | 0.79 |
| **Race (% White, non-Hispanic)** | 32% | 40% | 0.79 |
| **Hispanic ethnicity** | 19% | 19% | 1 |
| **Age, years (mean, range)** | 54 (22-77) | 54 (30-74) | 0.99* |
| **Donor type (% Deceased donor)** | 48% | 52% | 1 |
| **Induction type (% ATG)** | 26% | 35% | 0.58 |
| **CMV high risk (D+/R-)** | 10% | 0% | 0.24** |
| **CMV intermediate risk (R+)** | 87% | 81% | 0.73 |

\* p-value from t-test
\*\* p-value from Fisher's exact test

### Plasma proteome provides CMV-DNAemia specific protein profiles

To compare protein changes associated with CMV DNAemia, we analyzed the identified plasma protein data from CMV DNAemia positive patients and compared it with protein data from CMV DNAemia negative patients. A total of 241 proteins were used for this analysis (**Supplemental Table 1**). To analyze distinct proteomic features among patient CMV DNAemia status using a two-component linear discriminant analysis was used to distinguish clustering of patient CMV DNAemia status over time. LDA separated samples based on CMV DNAemia status among CMV DNAemia positive and CMV DNAemia negative cohort. Within CMV DNAemia positive cohort the samples were separated based on post-CMV DNAemia timepoint (**Figure 2A**). Among samples collected from CMV DNAemia individuals, samples collected 1 week and 1-month post-DNAemia are interspersed suggesting the signal of CMV infection persists until 1-month post-infection (**Figure 2A**).

**Figure 1:**
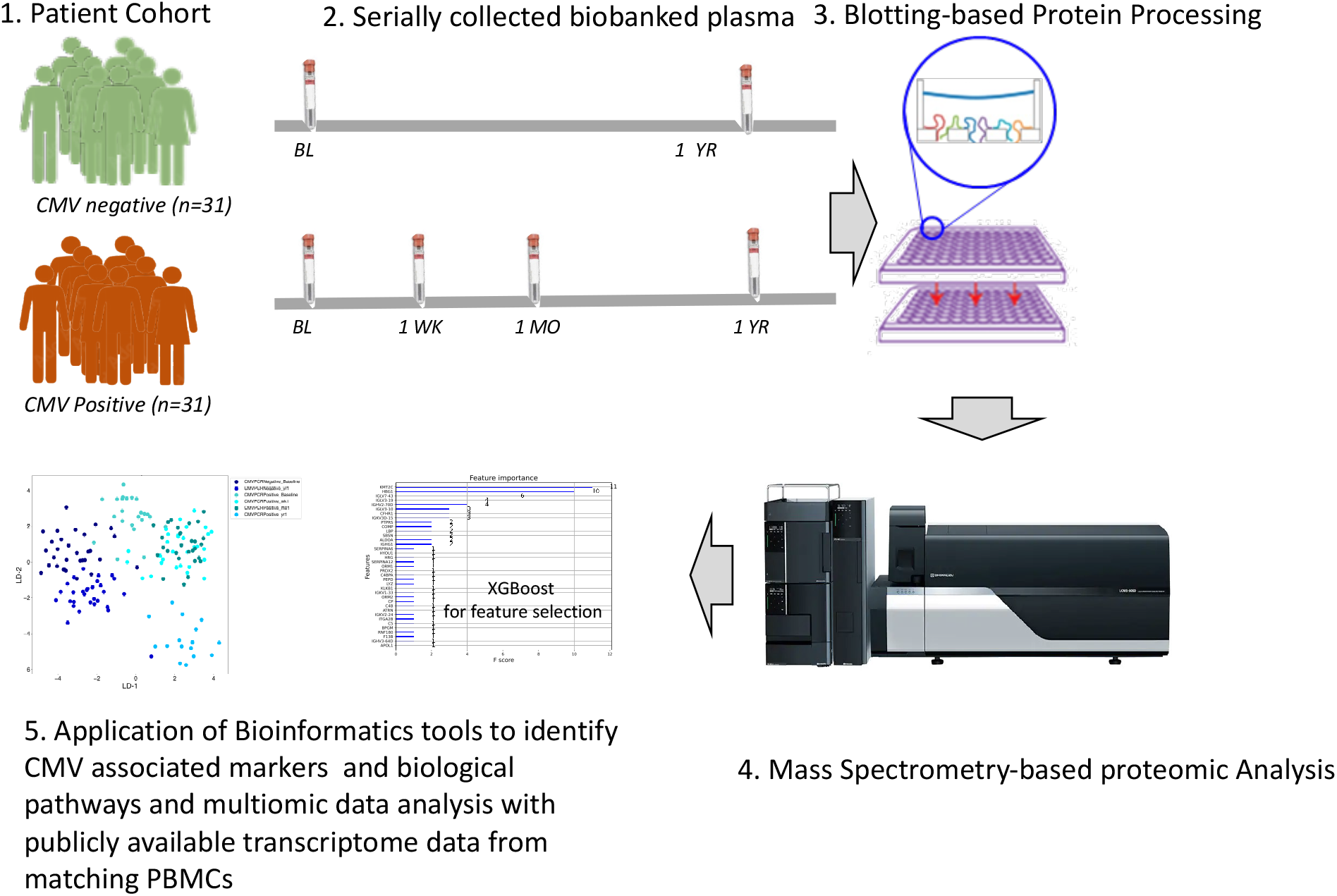
Study schematics. Proteomics analysis of plasma samples collected from kidney transplant recipients to identify CMV-associated proteins in kidney transplantation

**Figure 2:**
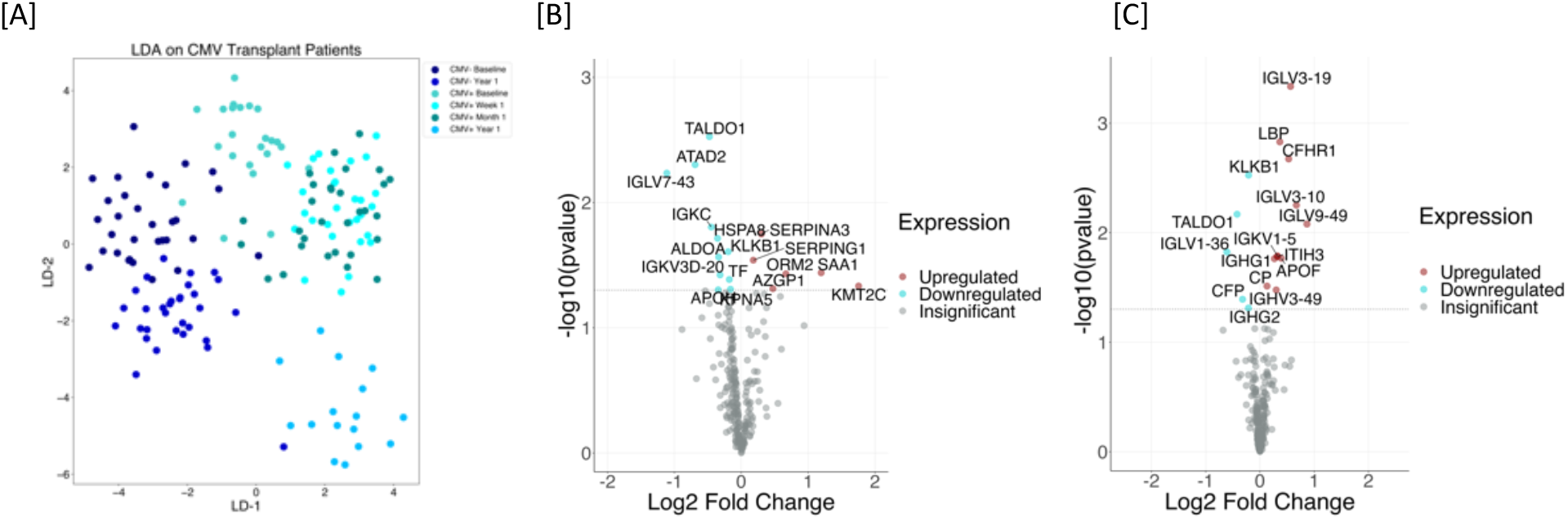
CMV DNAemia specific protein signatures exist pre-CMV DNAemia timepoint and post-CMV. **[A]** Plasma protein profiles for pre- and post-CMV DNAemia samples CMV DNAemia positive and negative cohort were analyzed using linear discriminant analysis (LDA). LDA was able to separate samples based on CMV status among CMV DNAemia positive and CMV DNAemia negative cohort. Within CMV DNAemia positive cohort the samples were separated based on post-CMV infection time. Among samples collected from CMV DNAemia positive individuals, samples collected 1-week and 1-month post-infection were interspersed suggesting the signal of CMV infection persists until 1-month post-infection. **[B]** Plasma samples were collected at pre-CMV infection time from CMV DNAemia positive and matching CMV DNAemia negative cohort. The analysis resulted in a set of 17 proteins whose levels were either increased (n=6) or decreased (n=11). The significance of the changes is demonstrated by a Volcano plot. The most significantly increased protein was Lysine Methyltransferase 2C (KMT2C) with 3.38 fold increase (p=0.05) in CMV DNAemia positive samples and most significantly decreased protein was Immunoglobulin Lambda Variable 7-43 (IGLV7-43) with 2.17 fold decrease (p=0.01). **[C]** We analyzed samples that were collected at 1-week and 1-month post-CMV infection from CMV DNAemia positive cohort and compared them against proteins profiles generated from CMV DNAemia negative cohort that included baseline and 1-year time-point. This analysis resulted in significant changes in 16 proteins with an increase in their level with CMV infection (n=11) and a decrease in their level with CMV DNAemia (n=5). Changes in individual proteins are presented as a volcano plot.

### Pre-CMV DNAemia perturbation in plasma proteome

To investigate if there are changes in protein profiles pre-CMV infection we analyzed plasma protein profiles generated from samples collected at pre-CMV infection time from CMV DNAemia positive and matching CMV DNAemia cohort as described in **Table 1**. The analysis resulted in a set of 17 proteins whose levels were either increased (n=6) or decreased (n=11) **Table 2**. The significance of the changes is demonstrated by Volcano plot in **Figure 2B**. Relative fold increase and decrease and statistical significance is presented in a bubble plot (**Supplemental Figure 1)**. The most significantly increased protein was Lysine Methyltransferase 2C (KMT2C) with 3.38-fold increase (p=0.05) in CMV DNAemia positive samples and most significantly decreased protein was Immunoglobulin Lambda Variable 7-43 (IGLV7-43) with 2.17 fold decrease (p=0.01). A complete list of proteins with fold change and p value is provided in **Table 2**. The top 3 biological processes enriched by the significant proteins were platelet degranulation (FDR, 4.83E-06), acute inflammatory response (FDR, 0.0018), blood coagulation (FDR, 0.0018). A complete list is provided in **Supplemental Table 2**.

**Table 2.** pre-CMV-DNAemia protein signature for CMV DNAemia.

| **S.No.** | **Gene Symbol** | **Protein Name** | **fold change** | **P value** |
| --- | --- | --- | --- | --- |
| 1 | KMT2C | lysine methyltransferase 2C | 3.38 | 4.64E-02 |
| 2 | SAA1 | serum amyloid A1 | 2.29 | 3.64E-02 |
| 3 | ORM2 | orosomucoid 2 | 1.59 | 3.71E-02 |
| 4 | AZGP1 | alpha-2-glycoprotein 1, zinc-binding | 1.39 | 4.87E-02 |
| 5 | SERPINA3 | serpin family A member 3 | 1.23 | 1.78E-02 |
| 6 | SERPING1 | serpin family G member 1 | 1.13 | 2.88E-02 |
| 7 | KPNA5 | karyopherin subunit alpha 5 | 0.89 | 4.92E-02 |
| 8 | TF | transferrin | 0.88 | 4.11E-02 |
| 9 | KLKB1 | kallikrein B1 | 0.87 | 2.47E-02 |
| 10 | IGKV3D-20 | immunoglobulin kappa variable 3D-20 | 0.80 | 3.78E-02 |
| 11 | ALDOA | aldolase, fructose-bisphosphate A | 0.79 | 2.72E-02 |
| 12 | APOH | apolipoprotein H | 0.79 | 4.97E-02 |
| 13 | HSPA8 | heat shock protein family A (Hsp70) member 8 | 0.78 | 1.93E-02 |
| 14 | IGKC | immunoglobulin kappa constant | 0.73 | 1.57E-02 |
| 15 | TALDO1 | transaldolase 1 | 0.72 | 2.98E-03 |
| 16 | ATAD2 | ATPase family AAA domain containing 2 | 0.62 | 4.99E-03 |
| 17 | IGLV7-43 | immunoglobulin lambda variable 7-43 | 0.46 | 5.82E-03 |

### CMV DNAemia specific protein signature

**T**he LDA analysis demonstrates that the plasma proteomic profile of patients at 1-week and 1-month post-CMV infection are similar and interspersed (**Figure 2A**). Next, we compared plasma protein profiles for samples that were collected at 1-week and 1-month post-CMV DNAemia from CMV DNAemia positive cohort and compared them against proteins profiles generated from CMV DNAemia negative cohort that included Pre-DNAemia and 1-year timepoint. Analysis resulted in significant changes in 16 proteins with an increase in their level with CMV infection (n=11) and a decrease in their level with CMV infection (n=5) (**Table 3**). Among the increased proteins mostly were immunoglobin complex components, IGLV9-49, IGLV3-10, IGLV3-19, IGKV1-5, IGHV3-49, IGHG1. Other were plasma proteins complement factor H related 1 (CFHR1), apolipoprotein F (APOF), Lipopolysaccharide Binding Protein (LBP), inter-alpha-trypsin inhibitor heavy chain 3 (ITIH3), and ceruloplasmin (CP), decreased proteins were immune complex components, IGHG2, IGLV1-36, complement factor P (CFP), kallikrein B1 (KLKB1), CFP (complement factor properdin), and transaldolase (TALDO1). Changes in individual proteins are presented as a Volcano Plot (**Figure 2C**). This demonstrated that there is a rise in the level of a set of plasma proteins immediately after CMV DNAemia.

**Table 3.** List of plasma proteins increased or decreased specific to CMV DNAemia (cross-sectional analysis).

|  |  |  | Fold Change | p value |
| --- | --- | --- | --- | --- |
| 1 | IGLV9-49 | immunoglobulin lambda variable 9-49 | 1.82 | 8.00E-03 |
| 2 | IGLV3-10 | immunoglobulin lambda variable 3-10 | 1.59 | 6.00E-03 |
| 3 | IGLV3-19 | immunoglobulin lambda variable 3-19 | 1.48 | 5.00E-04 |
| 4 | CFHR1 | Complement Factor H Related 1 | 1.44 | 2.00E-03 |
| 5 | APOF | Apolipoprotein F | 1.31 | 2.70E-02 |
| 6 | LBP | Lipopolysaccharide Binding Protein | 1.29 | 1.50E-03 |
| 7 | IGKV1-5 | Immunoglobulin Kappa Variable 1-5 | 1.25 | 1.60E-02 |
| 8 | ITIH3 | Inter-Alpha-Trypsin Inhibitor Heavy Chain 3 | 1.25 | 1.70E-02 |
| 9 | IGHV3-49 | Immunoglobulin Kappa Variable 3-49 | 1.23 | 3.30E-02 |
| 10 | IGHG1 | Immunoglobulin Heavy Constant Gamma 1 | 1.20 | 1.70E-02 |
| 11 | CP | ceruloplasmin | 1.09 | 3.00E-02 |
| 12 | KLKB1 | Kallikrein B1 | 0.86 | 3.00E-03 |
| 13 | IGHG2 | Immunoglobulin Heavy Constant Gamma 2 | 0.86 | 5.00E-02 |
| 14 | CFP | Complement Factor Properdin | 0.80 | 4.00E-02 |
| 15 | TALDO1 | Transaldolase 1 | 0.75 | 8.00E-03 |
| 16 | IGLV1-36 | immunoglobulin lambda variable 1-36 | 0.66 | 1.50E-02 |

We also analyzed proteomics data to see long-term proteomic changes 1 year after DNAemia by comparing protein profiles of 12-month post-DNAemia time points in between positive and negative DNAemia cohorts. As expected, the signature of CMV at the time of DNAemia was lost.

We identified only 4 proteins, namely, IGLV3-19 (1.7-fold increase, p =0.003), IGHG3 (1.5 fold increase, p=0.02), PSMB4 (1.3 fold increase, p=0.03), CFHR5 (1.8 fold increase, p=0.03) were increased in CMV DNAemia positive cohort indicating a long-term residual impact of CMV DNAemia in kidney transplant recipients (**Supplemental Table 3**).

### Temporal Proteome changes due to CMV DNAemia

We performed an analysis of the proteomic dataset for quantitative alteration of plasma proteins that correspond to CMV DNAemia. Protein profiles of baseline samples of CMV DNAemia positive patients were compared with protein profiles of 1-week post-infection samples which resulted in ten significantly changed proteins. The significant proteins expression direction and significance is presented with a volcano plot (**Figure 3A**). Increased proteins at the time of CMV DNAemia positive infection (1-week post DNAemia) included Serpin Family A Member 12 (SERPINA12) with p value 0.01 and 2.47 fold increase and Immunoglobulin Heavy Variable 3-72 (IGHV3-72) with p value 0.02 and 1.65 fold increase. Transthyretin (TTR) and Lysine Methyltransferase 2C (KMT2C) (**Table 4**). The proteins were involved in enrichment of biological processes such as complement activation (FDR=0.03), Humoral immune response (FDR=0.01), innate immune response (FDR=0.01) (**Supplemental Table 4**).

**Table 4.** List of plasma proteins increased or decreased on longitudinally collected CMV DNAemia samples (temporal analysis).

| S. No. | Symbol | Protein Name | Fold Change | p value |
| --- | --- | --- | --- | --- |
| 1 | LBP | Lipopolysaccharide Binding Protein | 1.44 | 2.96E-03 |
| 2 | SERPINA12 | Serpin Family A Member 12 | 2.47 | 6.92E-03 |
| 3 | PGLYRP1 | Peptidoglycan Recognition Protein 1 | 1.43 | 1.55E-02 |
| 4 | CP | Ceruloplasmin | 1.22 | 1.76E-02 |
| 5 | IGHV3-72 | Immunoglobulin Heavy Variable 3-72 | 1.65 | 2.23E-02 |
| 6 | C8A | Complement C8 Alpha Chain | 1.19 | 2.30E-02 |
| 7 | TTR | Transthyretin | 0.72 | 3.93E-02 |
| 8 | C4BPA | Complement Component 4 Binding Protein Alpha | 1.16 | 4.20E-02 |
| 9 | CFH | Complement Factor H | 1.11 | 4.25E-02 |
| 10 | KMT2C | Lysine Methyltransferase 2C | 0.23 | 4.73E-02 |

**Figure 3:**
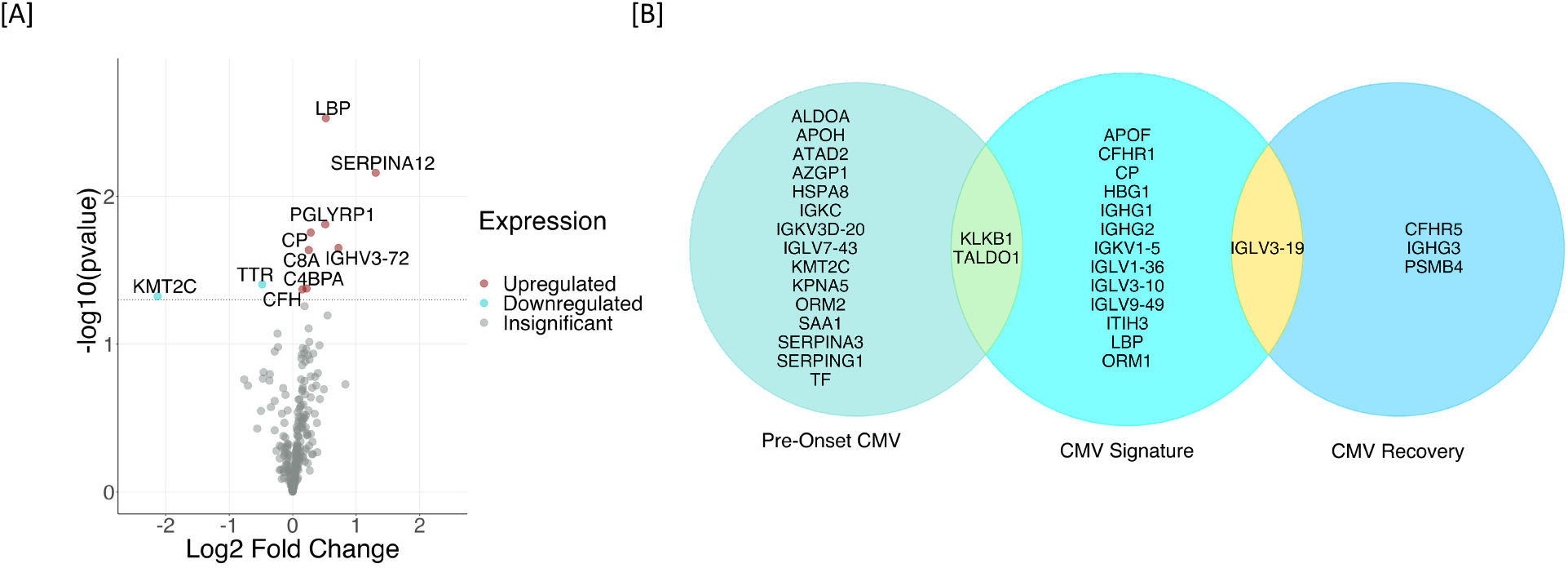
Temporal Proteome changes due to CMV DNAemia. **[A]** Protein profiles of baseline samples of CMV DNAemia positive patients were compared with protein profiles of 1-week post-infection samples which resulted in ten significantly changed proteins. **[B]** CMV DNAemia induced plasma proteomic profile overtime was analyzed. A summary of protein changes due to CMV DNAemia is presented for three timepoints namely pre-CMV DNAemia, immediately post-CMV DNAemia, and at recovery or 1-yr post CMV DNAemia.

**Figure 4:**
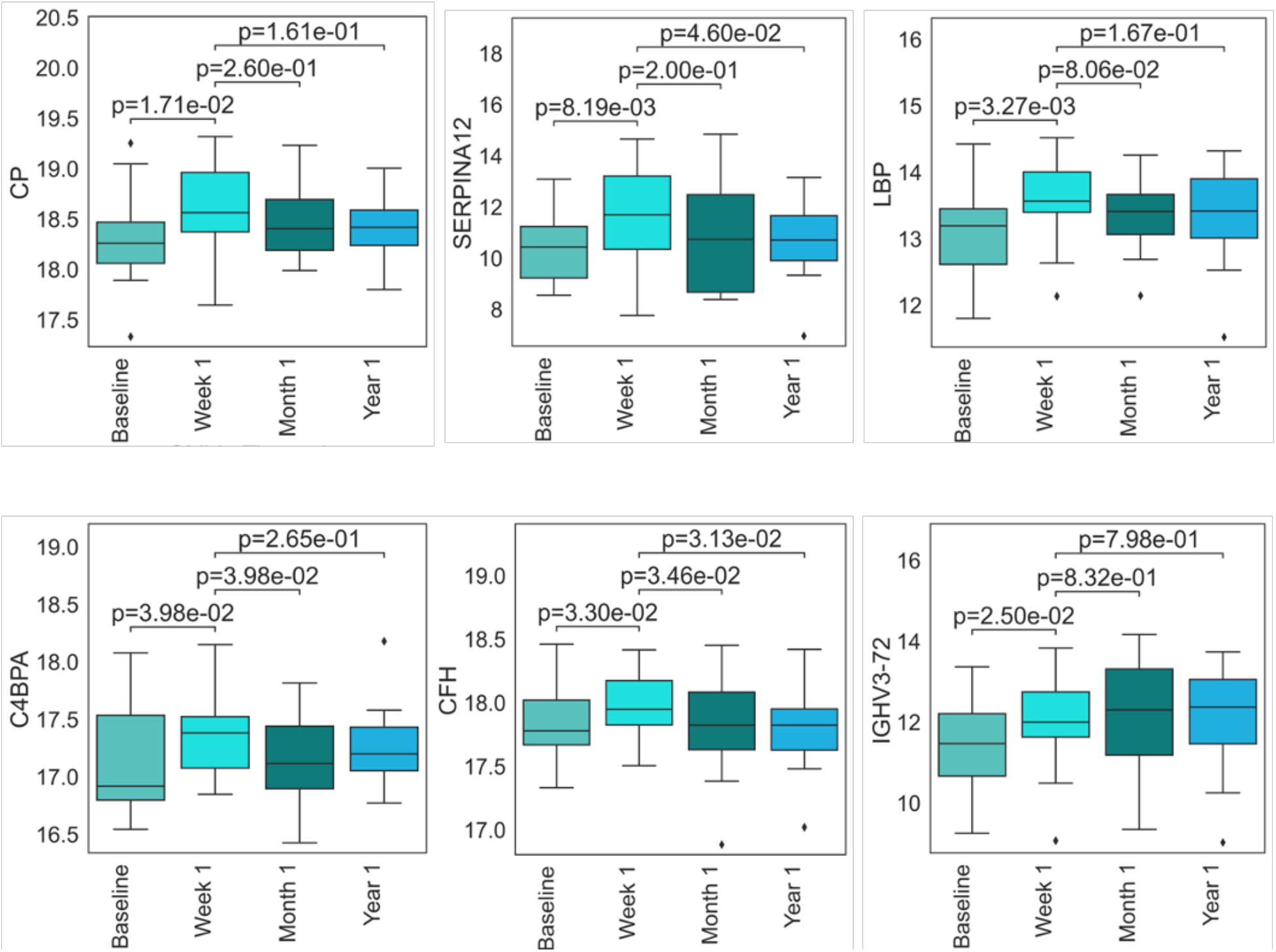
CMV DNAemia-specific proteins show increase in their level in 1-week post-DNAemia. Protein profiles immediately after CMV DNAemia were enriched with ceruloplasmin (CP), vaspin (SERPINA12), Lipopolysaccharide-binding protein (LBP), C4b-binding protein alpha chain (C4BPA), Complement Factor H (CFH), immune complex protein (IGHV3-72).

### CMV DNAemia induced proteomic signatures

A summary of protein changes due to CMV DNAemia is presented in **Figure 3B** listing plasma proteomic differences in the CMV DNAemic cohort pre-, immediately post- and 1 year post -DNAemia. Pre-CMV DNAemia samples showed an increase in serine protease inhibitors SERPINA3 and SERPING1 and enrichment of proteins involved in acute inflammatory responses such as Serum amyloid protein A (SAA1). Protein profiles immediately after CMV DNAemia showed higher amounts of immunoglobins and immunoglobulin complex components; ceruloplasmin (CP), vaspin (SERPINA12), Lipopolysaccharide-binding protein (LBP), C4b-binding protein alpha chain (C4BPA), Complement Factor H (CFH) and immune complex protein (IGHV3-72) were significantly increased in 1-week post-CMV DNAemia. The increased levels of these proteins were significantly lower both 1-mo- and 1-yr-post-DNAemia plasma samples (**Figure 5**).

Using XGBoost, a decision-tree-based ensemble machine learning algorithm that uses a gradient boosting framework, we developed a classifier that provided an accuracy of 88%, sensitivity of 85%, specificity of 91%, and a ROC AUC of 96% in identifying patients with CMV DNAemia. This classifier also identified a subset of 37 proteins **(Table 5)** that contributed to distinguishing CMV DNAemia **(Figure 5)**.

**Table 5.** Plasma protein signature for identification of CMV DNAemia positive patients in kidney transplant cohort.

| Gene Symbol | Protein Name | Impact Score (F-Score) |
| --- | --- | --- |
| ALDOA | aldolase, fructose-bisphosphate A | 11 |
| APOL1 | apolipoprotein L1 | 10 |
| ATRN | attractin | 6 |
| BPGM | bisphosphoglycerate mutase | 4 |
| C4B | complement C4B (Chido blood group) | 4 |
| C4BPA | complement component 4 binding protein alpha | 3 |
| C5 | complement C5 | 3 |
| CFHR1 | complement factor H related 1 | 3 |
| COMP | cartilage oligomeric matrix protein | 2 |
| CP | ceruloplasmin | 2 |
| F13B | coagulation factor XIII B chain | 2 |
| HRG | histidine rich glycoprotein | 2 |
| HYOU1 | hypoxia up-regulated 1 | 2 |
| IGHG1 | immunoglobulin heavy constant gamma 1 (G1m marker) | 2 |
| IGHV2-70D | immunoglobulin heavy variable 2-70D | 1 |
| IGHV3-64D | immunoglobulin heavy variable 3-64D | 1 |
| IGKV1-33 | immunoglobulin kappa variable 1-33 | 1 |
| IGKV2-24 | immunoglobulin kappa variable 2-24 | 1 |
| IGKV3D-15 | immunoglobulin kappa variable 3D-15 | 1 |
| IGLV3-10 | immunoglobulin lambda variable 3-10 | 1 |
| IGLV3-19 | immunoglobulin lambda variable 3-19 | 1 |
| IGLV7-43 | immunoglobulin lambda variable 7-43 | 1 |
| ITGA2B | integrin subunit alpha 2b | 1 |
| KLKB1 | kallikrein B1 | 1 |
| KMT2C | lysine methyltransferase 2C | 1 |
| LBP | lipopolysaccharide binding protein | 1 |
| LYZ | lysozyme | 1 |
| ORM1 | orosomucoid 1 | 1 |
| ORM2 | orosomucoid 2 | 1 |
| PEPD | peptidase D | 1 |
| PRDX2 | peroxiredoxin 2 | 1 |
| PTPRS | protein tyrosine phosphatase receptor type S | 1 |
| RNF180 | ring finger protein 180 | 1 |
| SBSN | suprabasin | 1 |
| SERPINA12 | serpin family A member 12 | 1 |
| SERPINA6 | serpin family A member 6 | 1 |
| HGB1 | Heterotrimeric G-protein beta subunit | 1 |

**Figure 5:**
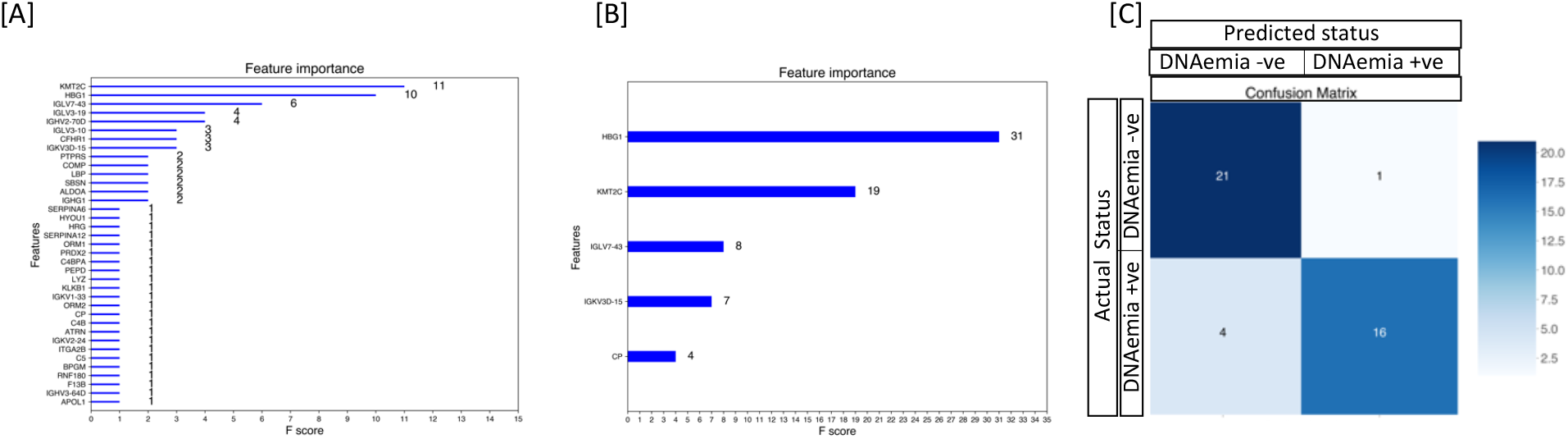
CMV DNAemia specific protein signatures exist pre-CMV DNAemia timepoint and post-CMV. **[A]** Using the protein profiles of DNAemia positive and DNAemia negative patients, we identified a classifier that provided a sensitivity of 85% and specificity of 91% in identifying CMV Dnaemia with a ROC AUC of 0.96. This classifier also identified a subset of 37 proteins that contributed with higher significance in distinguishing CMV DNAemia positive from CMV DNAemia negative samples **[B]** XGBoost-Boruta algorithm was applied to detect the top essential protein features. This process reduced the 241-protein list down to 5 proteins and increased AUC by 1%. This set of proteins included HBG1, KMT2C, IGLV7-43, IGKV3D-15, and CP **[C]** XGBoost machine learning was used on 70% samples to identify CMV DNAemia specific protein signature of 5 proteins. When tested for its performance on 41 samples (30%) with 21 CMV DNAemia positive and 20 CMV DNAemia negative status. The validation of the panel of proteins to identify CMV DNAmeia positive status irrespective of post CMV DNAemia time distinguished CMV positives with high accuracy of 88% and 95% specificity.

Furthermore, we used feature selection with the Boruta algorithm.The algorithm ran at 800,000 iterations was integrated to detect the top essential protein features. This process reduced the 241-protein list down to 5 proteins and increased AUC by 1%. This set of proteins included HBG1, KMT2C, IGLV7-43, IGKV3D-15, and CP **(Figure 5B)**. When tested for its performance on 41 samples with 21 CMV DNAemia positive and 20 CMV DNAemia negative status. The validation of the panel of proteins to identify CMV DNAmeia positive status irrespective of post CMV DNAemia time distinguished CMV positives with sensitivity at 80%, specificity at 95%, **(Figure 5C)** along with a ROC AUC of 0.97.

### Comparing plasma proteome with the transcriptome of PBMCs in matching blood samples

We performed a comparative multiome analysis of proteomics data from the plasma samples and transcriptome data available from the matching PBMCs from the same cohort which has been analyzed separately in a previously published paper^6^. Given two different milieus interrogated by transcriptomics (PBMCs) and proteomics (plasma) a significant overlap of target genes/proteins were not expected. Across all gene expression data and protein data, the average correlation for the most significant targets at the time of CMV DNAemia was 0.11, with the highest observed correlation between proteome and transcriptome was seen in immunoglobulin heavy constant gamma 1 (IGHG1) (Pearson, r=0.44) (**Supplemental Fig 3A**). There are six unique readouts when observing the differentially expressed CMV DNAemia positive signature between transcriptome and proteome. Among the six gene and proteins existing in CMV signature, Orosomucoid 1 (ORM1) holds a positive correlation (Pearson, r=0.17) (**Supplemental Fig 3B**). ORM1 gene encodes a key acute phase plasma protein which is also known to be decreased in blood samples infected with SARS-CoV-2^26^.

## Discussion

CMV modulates host immunity through several sophisticated strategies to achieve persistence in infected individuals. One of the mechanisms of modulation is reported to be the deployment of proteins to target host factors for degradation and suppression on host immunity against the virus in an *in vitro* model of CMV infection^27^. The virus has an impact on host gene and protein expression^6,27^. We recently reported the establishment of subset of prememory-like NK cells expressing NKG2C and lacking Fc epsilon RIgamma increased during viremia in CMV viremic patients^28^. An interactome study of CMV-host and virus-virus protein interactions identifies multiple degradation hubs^13^. These virus-host protein interactions enable the virus to evade host-immunity^29^. These cited works use either peripheral blood cells or in vitro cell system to study CMV’s role in CMV disease in humans. We performed proteomic analysis of plasma samples using blotting methodology^4,23^ and made significant observations about plasma protein markers and pathways associated with CMV DNAemia. This is first of its kind of studies that has been carried out in plasma samples from kidney transplant patients with and without DNAemia.

Dimensionality reduction with LDA based on 241 proteins identified in plasma of transplant patients successfully differentiated patients with and without CMV DNAemia. Furthermore, LDA separated patients into were groups based on post-CMV DNAemia timepoint. This highlights the fact that CMV induces changes in plasma protein composition that is unique to CMV DNAemia when compared to CMV DNAemia negative cohort. In addition, we also observed that plasma samples collected 1 week and 1-month post-DNAemia share proteomic composition in plasma suggesting the signal of CMV infection persists until 1-month post-infection.

Next, we identified CMV-specific proteins and biological pathways in pre-DNAemia plasma. We identified a set of 17 proteins (6 increased and 11 decreased) in pre-CMV DNAemia plasma samples which were enriched in biological processes such as platelet degranulation, acute inflammatory response, and blood coagulation. The most significantly increased protein was Lysine Methyltransferase 2C (KMT2C) is a potential tumor suppressor but has not been reported to be associated with CMV and is a novel finding^30^. Interestingly, expression of serum amyloid A1 (SAA1) protein was significantly elevated pre-DNAemia in patients that progressed to CMV

DNAemia. SAA is an acute phase protein with multiple immunological functions involved lipid metabolism and inflammation. In a cohort of lung transplant recipients, SAA concentrations were found to be significantly elevated in acute rejection and infection but not in stable transplants suggesting that SAA1 elevation is not an intrinsic response to graft^31^. Further evaluation of SAA1 levels in kidney transplant patients will provide validation of these results. These observations are intriguing and have the potential to lead to early markers for susceptibility for CMV infection among kidney transplant cohort. Earlier study profiling the transcriptional changes associated with CMV DNAemia did not detect any differences in pre-viremia signature of patients that went on to develop CMV DNAemia versus patients that did not^6^. This further underscore the importance of circulating proteins in the blood can provide crucial information about the immune status. Further validation of these proteins as early markers of CMV risk will provide better strategies in managing CMV infection in kidney transplantation.

It is known that CMV modulates host cells, downregulating and degrading hundreds of proteins^13^. Differential expression of proteins has been reported to control innate immunity linked to host response to viral infection^32^. Using in vitro system, previously published report listed 71 human proteins and 12 proteins encoded by known viral open reading frames (ORFs)^33^. For the first time, we identified plasma proteins whose level is significantly altered with CMV DNAemia. Increased immunoglobin complex components, IGLV9-49, IGLV3-10, IGLV3-19, IGKV1-5, IGHV3-49, IGHG1, plasma proteins such as complement factor H related 1 (CFHR1), apolipoprotein F (APOF), Lipopolysaccharide Binding Protein (LBP), inter-alpha-trypsin inhibitor heavy chain 3 (ITIH3), and ceruloplasmin (CP) provides information on perturbation on plasma proteomic landscape due to CMV DNAemia. Given, the different outcomes of CMV infection such as latent infection, subclinical infection, active infection, and CMV disease these plasma proteins offer potential biomarkers to stratify CMV infection types.

Despite, the study’s first-of-its-kind nature we acknowledge certain limitations. This study was done on samples collected from a single center. We do not know the impact of antiviral therapy on post DNAemia protein profiles and majority of patients have a history of CMV infection which may have contributed to plasma protein profile as reactivation of CMV instead of primary infection.

In summary, we believe the data presented provides insight on CMV DNAemia induced biological perturbations and potential protein markers for CMV infection. Further larger scale follow-up studies will further validate mechanisms and biomarkers for CMV disease and infection.

## Data Availability Statement

Proteomics data generated from this study has been deposited in IMMPORT database with accession number SDY2007, under workspace ID 6060 “Mapping Immune Responses to CMV in Renal Transplant Recipients, UCLA”. https://immport.niaid.nih.gov/research/study/studysearchmain#!/studysearch/viewStudyDetails/s tudySummarySDY2007

## Author Contributions

ER, MS, TS, JS, SB, DG, and MR designed the study. TS, MK, BF, HS generated the data. TS and PB drafted the manuscript. BF, TS and PB performed all data preprocessing and analyses. All authors contributed to the final analysis, data interpretation and manuscript writing and approved the submitted version.

